# Effects of Short-Term Breathwork on Respiration and Cognition

**DOI:** 10.1101/2025.11.17.688761

**Authors:** Lena Hehemann, Lisa Stetza, Christoph Kayser

## Abstract

Respiration is a unique physiological process that can operate automatically but can also be deliberately used to modulate the bodily state. Importantly, respiration is deeply intertwined with neural processes, influencing processes from arousal and attention to executive control. Although structured respiration (breathing practice) has been exploited for centuries, the immediate consequences of short periods of structured respiration for sensation and cognition remain poorly understood. The present study examined how brief, one-minute episodes of structured respiration affect respiration and performance in a subsequent sensory-cognitive task. Across two experiments, participants engaged in structured breathing practices manipulating either respiratory frequency (slow breathing vs. fast breathing) or the inhalation-exhalation ratio (short inspiration-long expiration; long inspiration-short expiration) prior to performing a visual emotion discrimination task. Immediately after breathing practice, participant’s respiration deviated from their baseline: each technique resulted in specific deviations of respiratory frequency, inhalation-exhalation ratio, or the occurrence of atypical respiratory cycles, suggesting technique-specific returns to regular respiration. During the subsequent emotion response accuracy or reaction times did not differ between breathing practices. However, we observed transient improvements in reaction times immediately following all practices, suggesting a brief facilitation of attentional or sensorimotor responsiveness following conscious breathing. Our findings indicate that even brief, consciously controlled respiration can transiently influence cognitive performance, highlighting the role of voluntary respiratory modulation in shaping brain function and behavior.

## Introduction

Respiration is more than a metabolic necessity; its rhythm is deeply embedded in the brain’s activity, influencing processes from arousal and attention to executive functions. While the effects of breathing practices (i.e. active breathwork) on well-being, stress and emotional regulation are well-documented (Chin et al. 2024; Fincham et al. 2023b; Fincham et al. 2023a; Kral et al. 2023; Röttger et al. 2021; Russo et al. 2017; Vanutelli et al. 2024; Zaccaro et al. 2018), their specific impact on cognitive performance has only recently received systematic attention. A growing body of research has demonstrated that during spontaneous, unstructured respiration performance in sensory-cognitive tasks systematically co-varies with the respiratory cycle. Participants tend to actively adapt their respiratory pattern to the temporal structure of laboratory tasks, with direct consequences for immediate task performance and subsequent memory performance (Brændholt et al. 2025; Flexmann et al. 1974; Grund et al. 2022; Huijbers et al. 2014; Perl et al. 2019). For example, across a large sample of laboratory tasks we have shown that both reaction times and the accuracy of perceptual reports co-vary systematically along the respiratory cycle, in a manner that renders the respiratory phase shortly prior to a response informative about the response speed and response accuracy (Harting et al. 2025; Johannknecht und Kayser 2022). These behavioral effects are paralleled by changes in large scale rhythmic brain activity, functional connectivity, and local markers of task-related attention in sensory and association cortices (Brændholt et al. 2025; Kluger et al. 2021; Liu et al. 2017; Maric et al. 2020; Perl et al. 2019; Stetza et al. 2025; Tort et al. 2020; Zelano et al. 2016) possibly driven by the extensive projections of brainstem respiratory centers to higher-order brain structures (Ito et al. 2014; Yackle et al. 2017)

This intricate connection between respiration, brain activity and cognitive performance provides a strong theoretical basis for how active breathwork could enhance cognitive performance. Indeed, one finds recommendations for structured or mindful respiration across social media, healthcare literature and much more. Controlled respiration encompasses various techniques that supposedly induce distinct physiological and psychological effects, both following short-term application and long-term practice. For example, practicing slow deep breathing has been shown to enhance parasympathetic tone and vagal activity (Pal et al. 2004), to foster sleep quality (Andas et al. 2023) and to improve general health markers and subjective ratings of well-being (Balban et al. 2023; Rahmathulla et al. 2024). In combination with simple exercise a 10-week deep breathing program was shown to reduce stress, mood disturbances, heart rate and cortisol levels (Perciavalle et al. 2017). But also short-term breathing practice can enhance autonomic function, mood, and cognition. Studies show that slow breathing increases heart rate variability, a physiological marker of reduced stress (De Couck et al. 2019), and promotes faster reaction times in a laboratory task (Gallego und Perruchet 1993). Hyperventilation or high-frequency yoga breathing, in contrast, was associated with immediate analgesic effects (Chalaye et al. 2009), enhanced emotional well-being (van Diest et al. 2014), increased cortical activity, attention-related processes (Telles et al. 2024; Budhi et al. 2024) and improved visual and auditory reaction times in a laboratory task (Neginhal et al. 2017).

Despite the general consensus that structured respiration exercises can have a positive influence on subjective wellbeing, physiological and cognitive functions, it remains unclear which specific respiratory patterns (fast, slow, or with altered inhalation-exhalation ratio) are most beneficial, and whether (or which) benefits can be obtained even after short periods of structured respiration. The possible pathways through which breathing practice could shape cognitive functions are multifold. Many practices induce changes in heart rate and its variability (De Couck et al. 2019; van Diest et al. 2014; Neginhal et al. 2017; Sroufe 1971). They can thereby alter vagal tone, which can enhance cognitive control, attentional regulation, and emotional stability via activation of the sympathetic nervous system (Smith et al. 2017; Thayer und Lane 2000; Thayer und Lane 2009). Studies also link breathing practices, such as slow breathing, to systematic changes in brain rhythms such as alpha band activity (Budhi et al. 2024; Telles et al. 2024).This is associated with attentional processes and arousal (Hanslmayr et al. 2011) suggesting that respiration-induced rhythmic neural activity may reflect one pathway through which voluntarily modulated respiration could directly influence sensory-cognitive processing. Moreover, diaphragmatic and deep breathing have been shown to reduce cortisol secretion (Ma et al. 2017; Martarelli et al. 2011; Perciavalle et al. 2017), underscoring their modulatory effect on the hypothalamic-pituitary-adrenal (HPA) axis, a key neuroendocrine system implicated in the regulation of cognitive functions (Erickson et al. 2003; Wolf 2003).

To date most studies investigating the effects of breathing practice have focused on one or two specific practices and examined their impact on a single temporal scale — either as brief interventions or long-term training. This limits comparability across studies and leaves open the question of whether any observed effects are specific to one particular breathing practice or reflect broader effects tied to active or conscious respiratory control in general, i.e. regardless of the specific pattern of respiration. In fact, it often remains unclear whether the observed behavioral benefits originate from directing attention towards the act of breathing itself, independent of the specific respiratory pattern, or whether they are linked to distinct temporal characteristics of the applied pattern.

We here ask how structured respiration for a period of one minute, which could easily be incorporated into daily activities, facilitates performance in a well-controlled sensory-laboratory task. We focus on an emotion discrimination task, for which previous work has shown that the accuracy by which participants perform the task systematically covaries with respiratory phase (Harting et al. 2025; Johannknecht und Kayser 2022; Zelano et al. 2016) possibly by systematically structuring neural activity in the limbic system (Zelano et al. 2016). Hence this may serve as a well-grounded model-task to better understand whether and which type of structured respiration can facilitate sensory-cognitive performance. Using this task, we compared four patterns of structured respiration: slow frequency breathing, fast frequency breathing, and breathing with long inhalation - slow exhalation periods (LISE), or with slow inhalation - long exhalation periods (SILE).

## Methods

A total of 65 participants took part in the study. All procedures were approved by the ethics committee of Bielefeld University and participants provided written informed consent prior to participation. All participants reported normal or corrected-to-normal vision. The data was collected anonymously, and it cannot be excluded that some individuals participated in both experiments described below. Participants were sampled from a typical student population and we expect their demographic data to be very similar to previous work (Park und Kayser 2021, 2022). While participants were informed about the general aim of the study - to explore the relation between breathing practice and cognitive performance - they were not specifically informed of how their specific responses or reaction times would be analyzed.

The experiments were conducted in acoustically shielded booths (E: Box; Desone, Germany). Respiratory data was measured using a thermistor (Littlfuse Thermistor No. GT102B1K, Mouser electronics) inserted in a disposable medical oxygen mask, which covered the participants’ mouth and nostrils, as described previously (Harting et al. 2025; Johannknecht und Kayser 2022; Stetza et al. 2025). The thermistor captured temperature fluctuations associated with inhalation and exhalation. The voltage drop across the thermistor was amplified in a custom-made circuit and recorded via ActiveTwo EEG system (BioSemi BV) with a sampling rate of 1024Hz. Visual stimuli and task instructions were presented via a projector using MATLAB (v. R2017a; The MathWorks, Inc., Natick, MA) and the Psychophysics Toolbox (v. 3.0.18).

### Experimental Paradigm

The data were obtained in two separate experiments employing two very similar variations of an emotion discrimination task derived from previous work (Johannknecht und Kayser 2022; Harting et al. 2025). Each trial started with a fixation period (0.7-1 s, uniform) followed by the presentation of a face showing one of two emotional expressions (duration of image presentation 0.16s, covering 11.8 degrees of visual angle). The images were taken from the FACES-database (Ebner et al. 2010). For the experiment we selected for 100 individuals two images depicting anger and two depicting disgust. Within a given block, each image was presented only once, ensuring no repeated exposure within blocks. Participants responded using the left or right arrow key on a computer keyboard. Inter-trial intervals consisted of a fixed component of 1200 ms and a random component of up to 300 ms. In Experiment 1, 45 participants classified the emotions anger and disgust across 3 blocks per technique of 133 trials each. In Experiment 2, 20 participants classified the emotions happiness and sadness across 2 blocks per technique of 100 trials each. Prior to the experiment, we recorded each participant’s spontaneous respiratory frequency for a duration of two minutes to establish an individual baseline. Prior to each block of the emotion task, and after half the trials of each block, participants performed one of the breathing techniques described below for 60 seconds.

### Respiratory training

Prior to the experimental session, participants received standardized verbal and visual instructions on each of the designated breathing techniques. To ensure proper execution, each technique was once practiced for one minute under the supervision of the experimenter. Participants were instructed to breathe exclusively through their nose during the structured breathing periods and were informed that they could resume their natural pattern immediately following the breathing practice. The breathing practice was guided by a continuous visual cue presented on a computer screen: a dynamically animated circle that expanded during the inhalation phase and contracted during the exhalation phase, providing a clear, time-synchronized pacing signal (min. visual angle: 1.5 degrees; max. visual angle: 5.4 degrees). During the actual experiment participants performed the breathing practice following these instructions but without direct supervision by the experimenter.

In experiment 1 we compared two breathing techniques manipulating the frequency of respiration: a slow breathing technique, implemented as box breathing, and a fast breathing technique. Box breathing consisted of four sequential phases of equal duration: inhalation, breath-hold, exhalation, and another breath-hold. To individualize the duration of each phase, participants completed a standardized breath-hold test prior to the breathing practice. This procedure was chosen based on evidence suggesting that breath-hold time is positively associated with vital lung capacity and serves as an indirect measure of peripheral chemoreflex sensitivity (Cherouveim et al. 2013; Trembach und Zabolotskikh 2017). Based on each participant’s breath-hold performance, the durations of each phase were set to either 3 seconds (breath hold time less than 20 seconds) or 5 seconds (breath hold time between 20 and 45 seconds), resulting in an overall respiratory cycle frequency of 0.08 Hz or 0.05 Hz, respectively. For the fast technique, the respiratory frequency was set at twice the participant’s baseline respiratory frequency. The respiratory cycle consisted of alternating inhalation and exhalation phases without periods of breath hold. Based on the definition for Tachypnea (increased or elevated respiratory frequency), doubling the spontaneous respiratory frequency should be sufficient to increase respiratory rate beyond the range covered by natural variation at rest, without at the same time mimicking very high frequency breathing that can be aversive.

In experiment 2 we compared three conditions: two conditions with altered inhalation to exhalation ratios (SILE and LISE) (Röttger et al. 2021; Strauss-Blasche et al. 2000) and a no-intervention condition (NOI). In SILE (*Short Inspiration - Long Expiration*), inspiration and expiration were paced following a 1:2 ratio, emphasizing prolonged expiration, while in LISE (*Long Inspiration - Short Expiration*), the ratio was inverted and emphasized prolonged inhalation. For both, the overall respiratory frequency was matched to each participant’s spontaneous respiratory frequency. The no intervention condition contained no specific practice and served as a baseline condition. For this, prior to the onset of the block, participants remained seated in the experimental room for one minute, engaging in spontaneous, self-paced breathing. As during blocks with actual breathing techniques, participants initiated the no intervention blocks by pressing a button. However, instead of a breathing exercise, they were presented with a 60 second idle time before the first trial.

### Preprocessing of respiratory data

The respiratory data were preprocessed similarly as in our previous work (Harting et al. 2025; Stetza et al. 2025). First, the data were filtered using a third-order Butterworth bandpass filter with cutoff frequencies of 0.03 Hz (high-pass) and 6 Hz (low-pass). The filtered data were then resampled to 100 Hz using the FieldTrip toolbox (fieldtrip-20231015). To facilitate inter-participant comparisons, the respiratory signal was normalized to z-scores. The Hilbert transform was applied to derive the analytic signal, from which local peaks were identified. Individual respiratory cycles were extracted from 7-second windows centered on each peak, retaining only those peaks for further analysis where the z-scored signal exceeded a threshold of z = 0.5. Inhalation phases were defined as segments with a continuous positive slope prior to each peak, whereby interruptions shorter than 500 ms were interpolated. Exhalation phases were similarly defined as phases with a continuous negative slope following each peak, again interpolating interruptions shorter than 500 ms. Atypical respiratory cycles were identified by computing mean-squared distances between individual cycles and the participant-specific centroid of all cycles. Cycles with distances exceeding three standard deviations (SD) from the centroid were classified as atypical. Atypical cycles may reflect breath-holds, sighs, or clearing of the throat.

**Figure 1:**
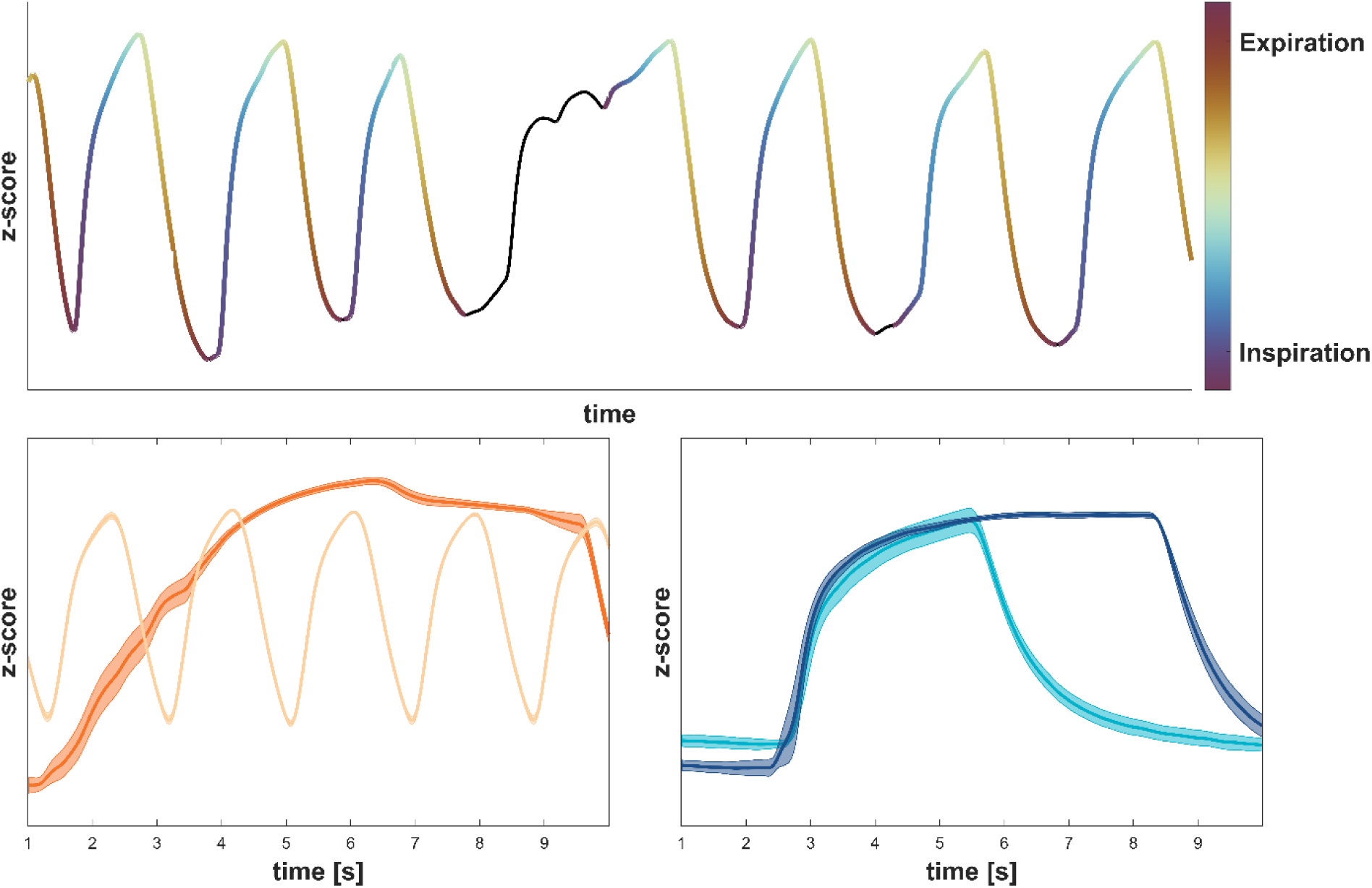
Exemplary respiratory traces illustrating spontaneous respiration and breathing practices. Each trace reflects the z-scored respiration signal over time. Upper panel: Illustration of the phase classification applied to the respiration signal. The color scale reflects the respiratory phase. Undefined segments are marked in black. Lower left panel: Representative traces for the Slow (Box Breathing, dark orange) and Fast (Hyperventilation, light orange) conditions from Experiment 1. Lower right panel: Representative traces for the SILE (light blue) and LISE (dark blue) conditions from Experiment 2.

### Processing of behavioral data

The behavioral data for the emotion task comprised trial-wise response accuracy and reaction times (RT). We removed a few individual outlier trials defined by the following criteria: trials were excluded if RT was shorter than 200 ms or exceeded three times the median absolute deviation from the participant’s individual RT distribution. This led to an exclusion of 4.91% ± 4.09% (mean ± std) of trials in experiment 1 and 5.29% ± 3.34% (mean ± std) of trials in experiment 2. For subsequent analysis RTs were square root transformed and normalized within participants by subtracting the average, thereby controlling for individual differences in overall response speed. For the analysis of RTs we focused only on trials with correct responses to isolate effects attributable to cognitive processing rather than motor errors or guessing. To quantify whether RTs follow a specific temporal trend following respiration practice, we binned these in groups of 10 trials for each participant. We then used a linear regression to derive the slope of the binned RTs against the time passed in the experimental block.

### Statistical analysis

Our main analyses probed for systematic differences in task performance between respiration techniques across participants. These tests were implemented separately for RTs and the fraction of correct responses (FCRs) using paired-sample *t*-tests. For each comparison we report *p*-values, Cohens d’ as a measure of effect size and Bayes factors (BF₁₀) to quantify evidence for the alternative hypothesis that the mean difference deviates from zero. The significance level was set to α = 0.05. To control for multiple comparisons within the same experiment, *p*-values were adjusted using the False Discovery Rate correction method (FDR) as proposed by (Benjamini und Yekutieli 2001). Note that p-values reported through the manuscript are the corrected p-values, and hence can in principle exceed 1.

To investigate potential temporal dynamics in RTs following breathing practice, we used a permutation test. We compared the actual slope of the binned RTs against time passed in the experiment with a surrogate distribution of slopes obtained after randomly shuffling the binned RTs and calculating the slopes in 2000 permutations. From the surrogate distribution we obtained the percentile p-value of the original slope.

## Results

### Respiration during the practice period

Across two experiments participants completed four types of breathing practice during one-minute breathing practice blocks prior to performing an emotion discrimination task. To characterize how participants breathed during the practice periods, we calculated their respiratory frequency and the inhalation-to-exhalation ratio and compared these to their baseline data (Figure 2).

**Figure 2.**
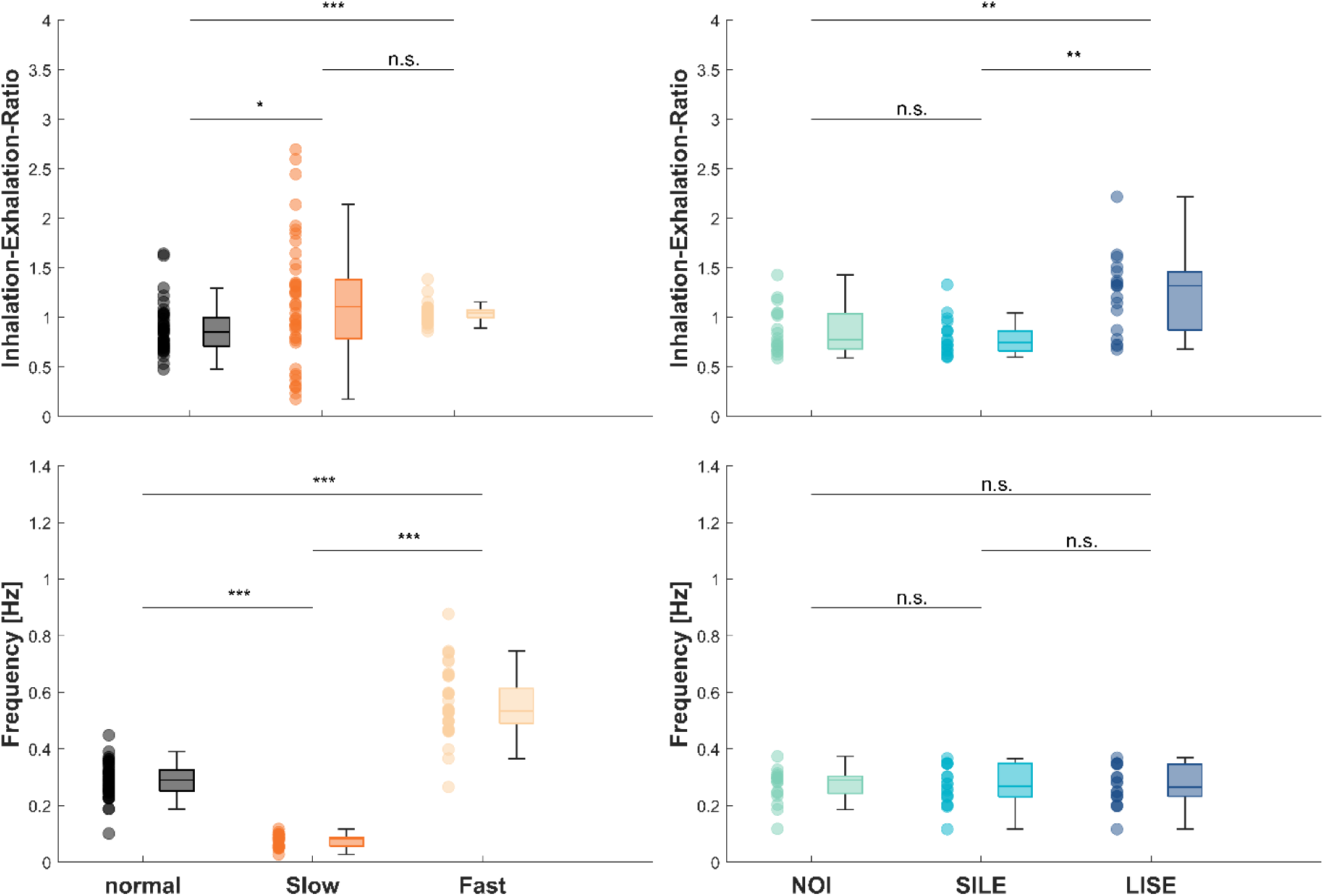
Respiratory characteristics during breathing practice for Experiment 1 (left panels) and Experiment 2 (right panels). The upper graphs show the comparison of inhalation-to-exhalation (I/E) ratio, the bottom graph those for respiratory frequency during the one-minute breathing practice (Slow: slow paced breathing, Fast: fast paced breathing, SILE: short inspiration - long expiration, LISE: long inspiration - short expiration). Baseline values were obtained from either a baseline recording prior to the experiment (Experiment 1,‘normal’) or from a one minute no intervention condition (experiment 2,‘NOI’). Data are presented as boxplots with individual participant values presented as dots. Statistical significance was determined using paired-sample t-tests; p-values were adjusted for multiple comparisons using the false discovery rate (FDR). Asterisks indicate statistically significant differences: * *p* < .05; ** p<0.01; *** p>0.001; n.s. non-significant

In experiment 1 participants (n=45) followed either a slow or a fast breathing practice. During these periods we observed clear and expected differences in respiratory frequencies: slow breathing elicited markedly lower respiratory frequencies compared to fast breathing (p <10^-5^, BF₁₀ = 10^27^, Cohens d’ = 5.9). Compared to baseline, fast breathing resulted in higher (p<10^-5^, BF₁₀ = 10^25^, Cohens d’ =2.9) and slow breathing in lower frequencies (p<10^-5^, BF₁₀ = 10^22^, Cohens d’ = 4.6). As expected, the average inhalation-to-exhalation ratio did not differ significantly between slow and fast breathing (p=0.523, BF₁₀ = −3.6, Cohens d’ = 0.2), but both ratios differed significantly compared to the baseline (slow: p = 0.021, BF₁₀ =5.0, Cohens d’ = 0.6; fast: p = p<10^-5^, BF₁₀ = 499.1, Cohens d’ = 0.9).

In experiment 2 participants (n=20) followed breathing practices aimed at manipulating the inhalation-to-exhalation ratio. As expected, we found no significant differences in respiratory frequency between conditions (NOI vs. SILE: *p* = 1.141, BF₁₀ = –3.7, Cohens d’ = 0.1; NOI vs. LISE: *p* = 1.141, BF₁₀ = –3.8, Cohens d’ = 0.1; SILE vs. LISE: *p* = 1.141, BF₁₀ = –2.3, Cohens d’ =0). As expected the breathing practices differed in their inhalation-to-exhalation ratios: the LISE condition exhibited a significantly higher ratio compared to the SILE condition (*p* = 0.004, BF₁₀ = 51.7, Cohens d’ = 1.5). Also, the inhalation-to-exhalation ratio in the LISE condition was significantly higher compared to participants’ baseline ratio (p = 0.007, BF₁₀ = 16.7, Cohens d’ = 1.2). In contrast, no significant difference was observed between the SILE condition and baseline (*p* = 0.690, BF₁₀ = –3.0, Cohens d’ = 0.3).

This shows that participants were able to adhere to the structured respiration, resulting in marked differences in respiratory frequency or inhalation-to-exhalation ratio.

### Respiration following the practice period

Inspecting the individual respiratory traces revealed that participants exhibited highly variable respiratory patterns following the breathing practice blocks (examples are shown in figure 3). Although explicitly instructed to return to natural, self-paced breathing after the practice, the respiration data did not necessarily reveal such a return. We observed three main patterns across participants: some participants continued to breathe similar as during the breathing practice for a prolonged time, while others quickly reverted to a respiratory pattern as observed during their baseline. Moreover, some participants showed extended breath-holding, sighs or very atypical respiratory cycles that could not be classified based on the recorded signal. The boundaries between these post-practice patterns appeared fluid, with some participants gradually transitioning between them or displaying different respiratory responses when repeating the same breathing practice.

**Figure 3:**
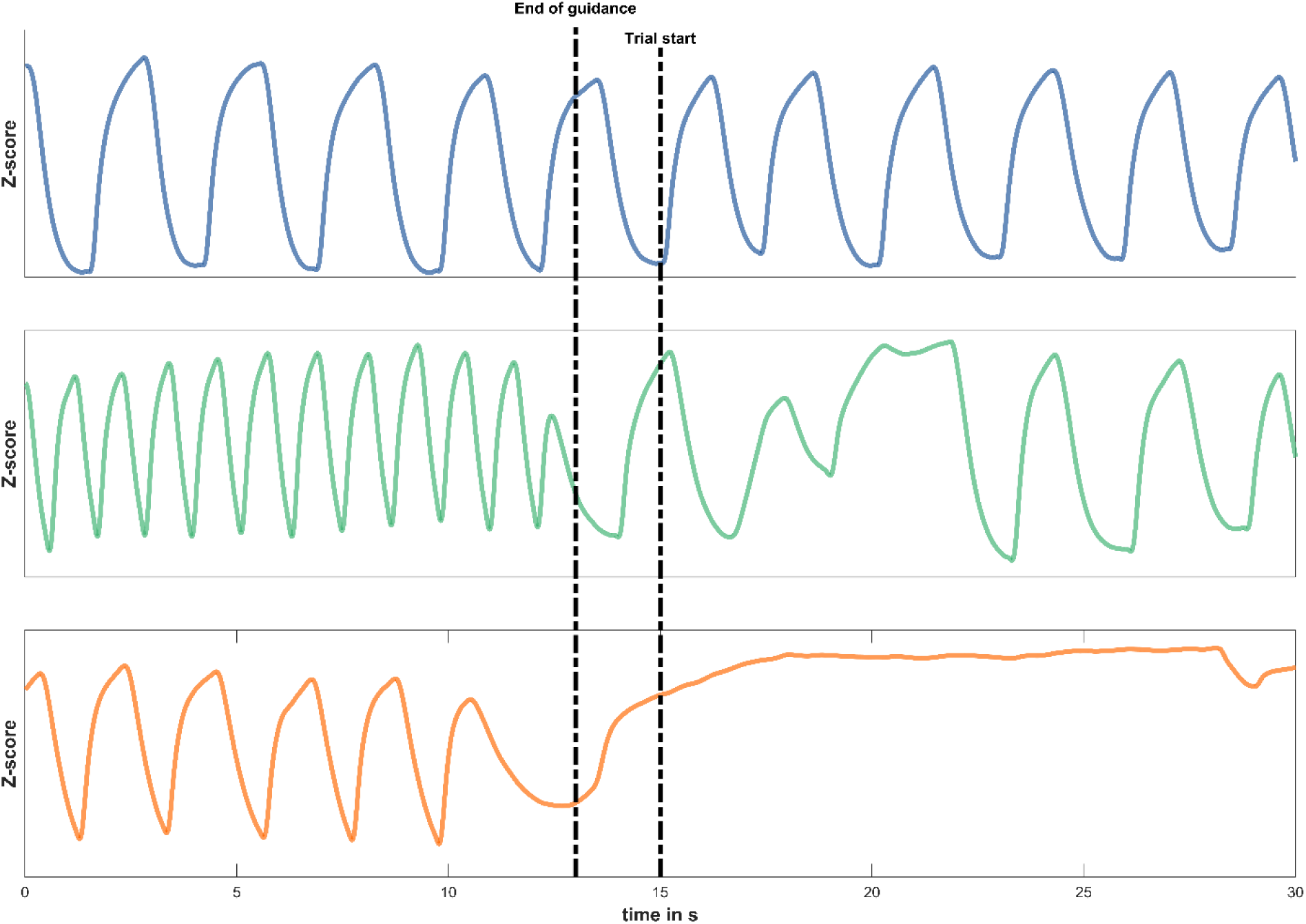
Exemplary respiratory traces from three participants illustrating individual differences in respiration following one-minute of fast breathing practice. Each trace displays the z-scored respiratory signal over time (in seconds), with the vertical black lines indicating the end of the guided breathing practice and task onset (2 seconds after the end of guided breathing practice). The top example continues to breath as during the breathing practice, the middle quickly returns to baseline and the bottom displays some forms of breath hold or extended sigh. Note that the frequency for hyperventilation was dependent on each participant’s baseline frequency.

To examine whether atypical respiratory patterns were associated with specific breathing practices, we calculated the proportion of data points that could not be assigned to a specific respiratory phase (inhalation or exhalation) within one minute following the end of guided breathing practice. Cycles containing a high amount of undefined data points may for example reflect breath-holds, sighs, or otherwise unstructured breathing. For remaining cycles, we calculated the frequency and the inhalation to exhalation ratio (see figure 4).

**Figure 4:**
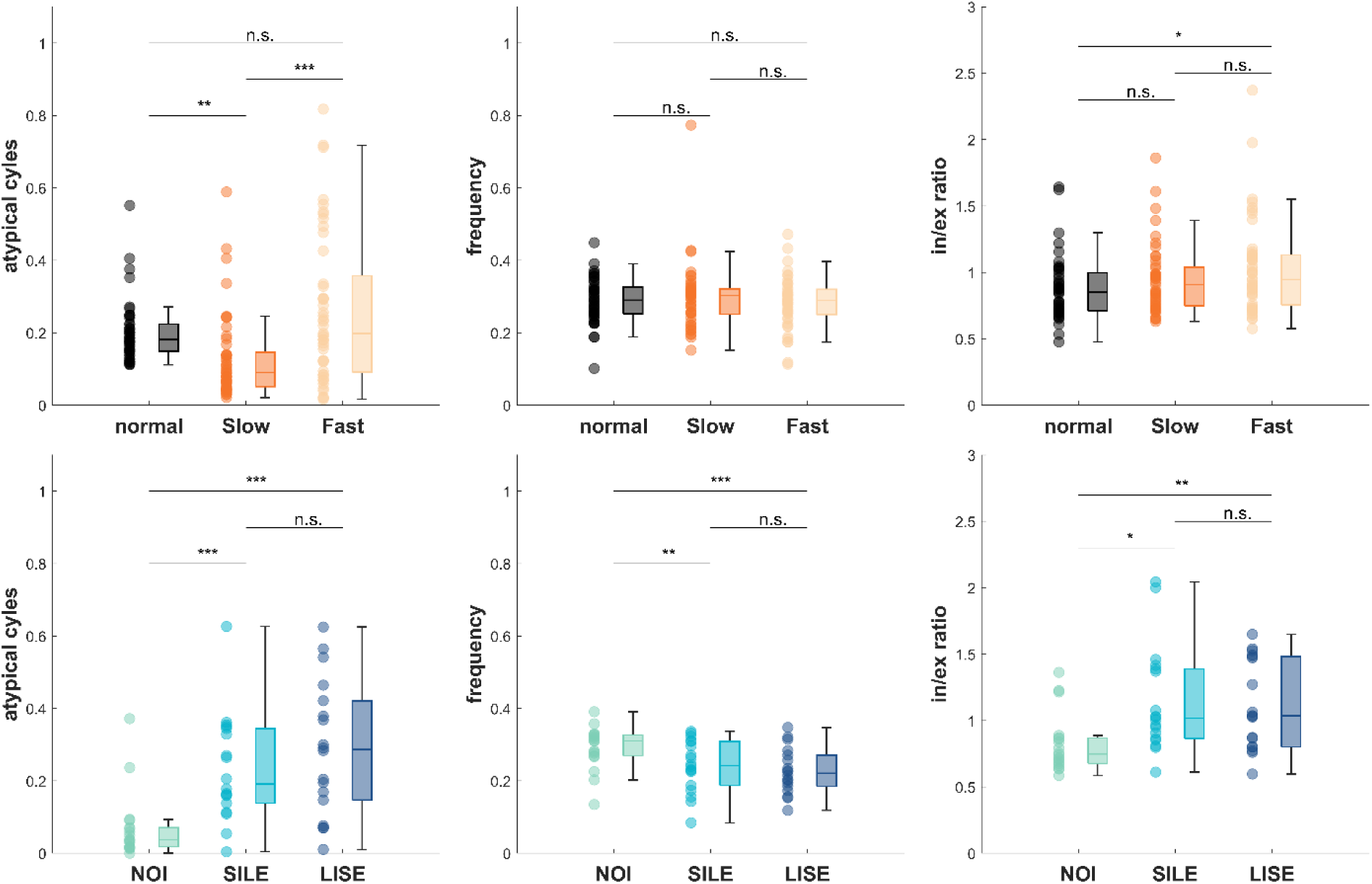
Respiration characteristics in a period of one minute following the breathing practice. The upper row presents results from Experiment 1 (Slow and Fast breathing), the bottom row from Experiment 2 (SILE, LISE, and NOI). The left panels show the percentage of undefined data points per participant (see Method section); the middle panels depict the respiratory frequency; the right panels illustrate the inhalation-to-exhalation ratio of those cycles that could be defined by the segmentation algorithm. Data is visualized using boxplots, with individual participant data shown as dots. Statistical comparisons were performed using paired-sample t-tests using the false discovery rate (FDR) to correct for multiple comparisons. Asterisks indicate statistically significant differences: * *p* < .05; ** p<0.01; *** p>0.001; n.s. non-significant.

In Experiment 1, the percentage of undefined data points differed significantly between the Fast and Slow breathing conditions (p <10^-5^, BF₁₀ = 4165.9, Cohens d’ = −0.8), with a higher proportion observed after Fast breathing. After the application of the slow breathing practice, the proportion of unclassified data points was reduced compared to the baseline condition (normal vs. Slow: p =0.001, BF₁₀=52.1, Cohens d’ =0.7). In contrast, the fast breathing practice elicited no statistically significant reduction or increase in undefined datapoints (normal vs. Fast: p=0.09, BF₁₀=1.0, Cohens d’ = - 0.38). There was no difference in respiratory frequency during the one-minute block after the breathing practice between conditions (Slow vs. Fast: p = 1.150, BF₁₀ = −4.5, Cohens d’ = 0.2; normal vs. Slow: p = 1.150, BF₁₀= −5.5, Cohens d’ = −0.1.; normal vs. Fast: p=1.15, BF₁₀=-5.5, Cohens d’ = 0.1). The inhalation-to-exhalation ratio differed between the recording of baseline respiration and the fast condition (p =0.041, BF₁₀ = 5.1, Cohens d’ = −0.5) but not between the recording of baseline respiration and the Slow condition (p = 0.095, BF₁₀=-1.0, Cohens d’ = −0.3) or between the breathing practice conditions (p=0.095, BF₁₀=-1.2, Cohens d’ = −0.3). (Figure 4, upper row)

In Experiment 2, we manipulated the inhalation/exhalation ratio. Significant differences emerged in the percentage of undefined data points after performing SILE and NOI (p = 0.001, BF₁₀ = 91.5, Cohens d’ = −1.3), as well as between LISE and NOI (p = 0.001, BF₁₀ = 141.8, Cohens d’ = −1.4). Respiratory frequency in the minute after both breathing practices differed from the no intervention condition (NOI vs. SILE: p = 0.003, BF₁₀ = 32.1, Cohens d’ = 0.7; NOI vs. LISE: p = 0.0003, BF₁₀ = 508.5, Cohens d’ = 1.0). Moreover, a significant difference in the inhalation/exhalation ratio was observed after SILE and LISE practice in comparison to the no intervention condition (NOI vs. SILE: p = 0.020, BF₁₀ =6.8, Cohens d’ = −1.0; NOI vs. LISE: p = 0.007; BF₁₀ = 32.2, Cohens d’ = −1.0). Statistically there was no difference between SILE and LISE condition, neither for the frequency (p = 0.406, BF₁₀=-2.1, Cohens d’ = 0.2) nor for the inhalation to exhalation ratio (p=1.412, BF₁₀ = −4.0, Cohens d’ = 0.1) in the minute after breathing practice. (Figure 4, lower row)

Overall, Fast, SILE, and LISE breathing practice exhibited a recovery of spontaneous respiration that differed from that observed following Slow respiration. Hence, slow breathing may specifically enable a fast and structured return to a spontaneous and regular respiratory pattern compared to the other techniques.

### Impact of respiration training on task performance

We analyzed the fraction of correct responses (FCRs, Figure 5) and reaction times (RTs, Figure 6) following each breathing practice. When considering the data from the entire experimental blocks we found no significant differences between breathing techniques in either Experiment 1 nor Experiment 2. The Bayes factors associated with individual comparisons were mostly in favor of no effect (indicated here by negative Bayes factors) but reflect either anecdotal or moderate evidence (Table 1).

**Figure 5:**
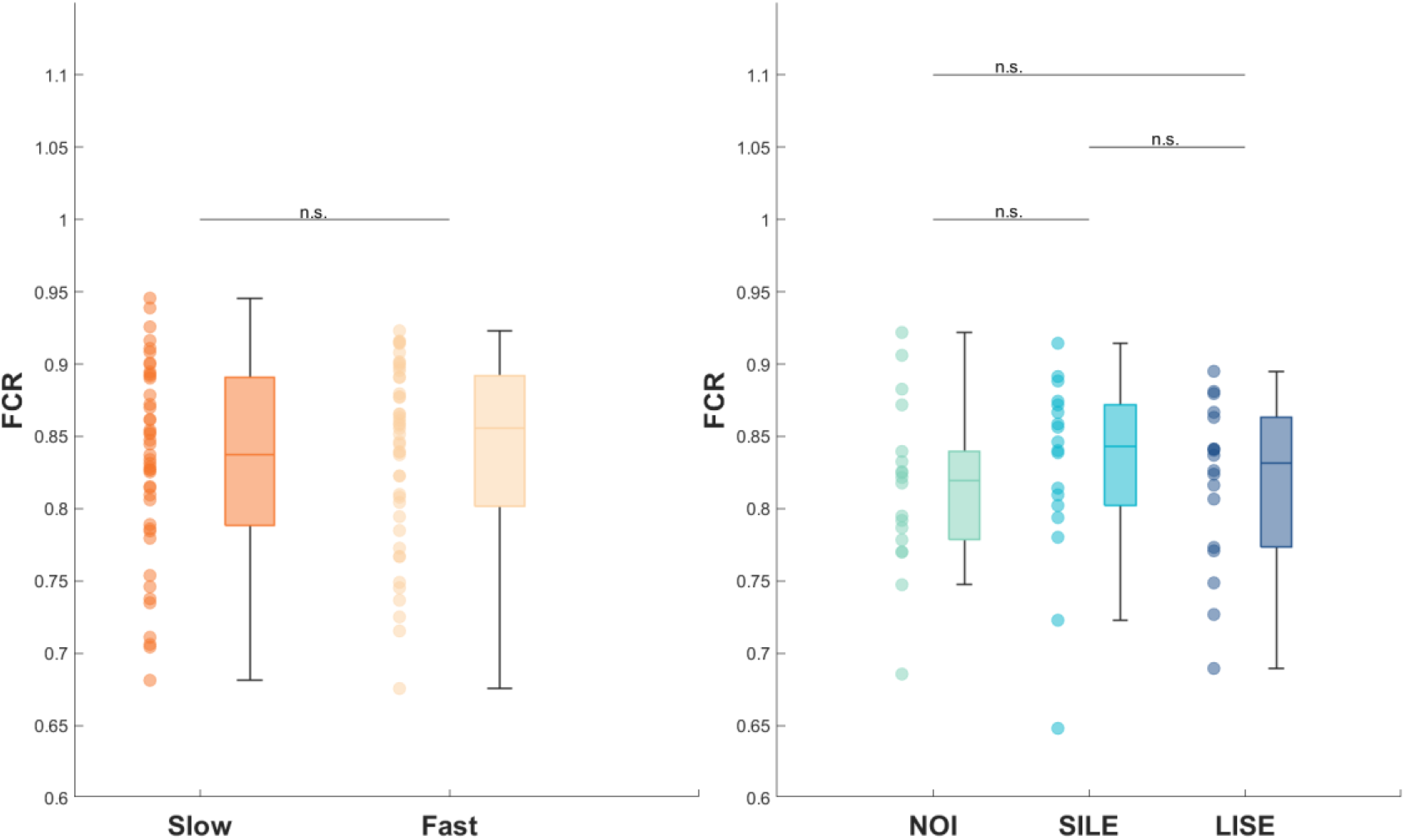
Response accuracy following breathing practice. The figure presents the average fraction of correct responses (FCRs) for each breathing practice. The left panel shows data from Experiment 1 (Slow and Fast breathing), the right panel shows data from Experiment 2 (NOI, LISE, SILE). The data is displayed as boxplots with individual data shown as dots. Pairwise statistical comparisons were performed using t-tests, with p-values corrected for multiple comparisons using the false discovery rate (FDR). Asterisks indicate statistically significant differences: * *p* < .05; ** p<0.01; *** p>0.001; n.s. non-significant.

**Figure 6:**
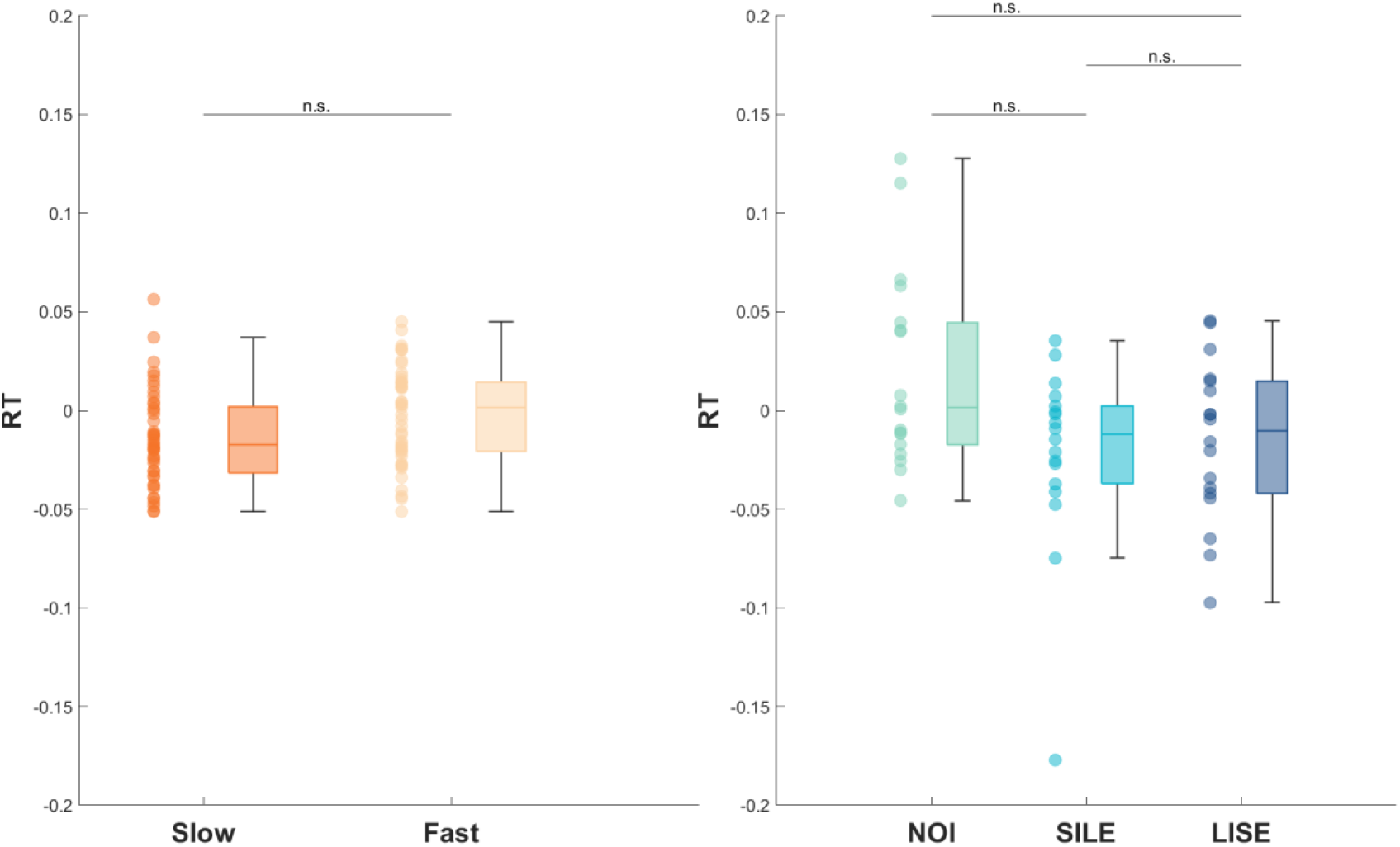
Reaction times following breathing practice. The figure shows the distribution of average reaction times (RTs) for each breathing condition. The left panel shows the results from Experiment 1 (Slow and Fast breathing), the right panel from Experiment 2 (NOI, LISE, SILE). For each participant, mean RTs were computed from all valid trials within the respective task block (see method section). Data is visualized using boxplots, with individual participant data shown as dots. Statistical comparisons between conditions were performed using paired-sample t-tests, and p-values were adjusted for multiple comparisons using the false discovery rate (FDR). Asterisks indicate statistically significant differences: * *p* < .05; ** p<0.01; *** p>0.001; n.s. non-significant.

**Tab 1:** Results of paired-sample t-tests comparing reaction times (RTs) and fraction of correct responses (FCRs) between breathing conditions in Experiment 1 and Experiment 2. Reported are p-values (corrected for multiple comparisons using the false discovery rate, FDR), together with BF₁₀ and Cohens d’ for each pairwise comparison.

|  | *FCR* | *RT* |
| --- | --- | --- |
| ***Slow >< Fast*** |  |  |
| *p-Value* | 0.199 | 0.095 |
| *BF<sub>10</sub>* | -2.8 | -1.6 |
| *Cohens d'* | 0.1 | 0.5 |
| ***NOI >< SILE*** |  |  |
| *p-Value* | 1.373 | 0.316 |
| *BF<sub>10</sub>* | -2.3 | 1.0 |
| *Cohens d'* | 0.2 | 0.8 |
| ***NOI >< LISE*** |  |  |
| *p-Value* | 1.373 | 0.316 |
| *BF<sub>10</sub>* | -3.3 | -1.3 |
| *Cohens d'* | 0.2 | 0.6 |
| ***SILE >< LISE*** |  |  |
| *p-Value* | 1.144 | 1.078 |
| *BF<sub>10</sub>* | -3.9 | -3.5 |
| *Cohens d'* | 0.1 | 0.2 |

While the overall effects of breathing practice seem to be small, visual inspection of the data revealed that there could be shorter-lived effects immediately following the practice. To quantify such effects, we binned the experimental trials into bins of ten trials. We then quantified whether RTs or FCR systematically change over the course of the experimental block using a regression analysis combined with a permutation test (Figure 7).

The results show that for all breathing practices the slopes of reaction times differed significantly from the null hypothesis of no significant slope. Notably, for all breathing practices these slopes were positive, indicating a progressive slowing of RTs after the application of breathing practice. Importantly, RTs during the NOI condition did not show a slope differing from 0, suggesting that this dynamic of reaction times (starting faster in a block and then slowing down) may be specific to having performed some form of breathing practice and not a general effect of fatigue during each block. (see Table2, Figure 7). To directly corroborate this conclusion, we compared the slopes between conditions in experiment 2. This revealed a significantly higher slope for SILE vs. the NOI condition (p = 0.015, BF₁₀= 3.7, Cohens d’ = −0.8) but not for LISE (p = 0.152, BF₁₀= −1.6, Cohens d’ = −0.4).

For the fraction of correct responses (FCR) this analysis revealed a significant negative slope (percentile in random permutation test: 0.13; slope: −0.011) for the fast breathing condition, indicating a decline in accuracy over time. In contrast, for all other breathing techniques as well as the NOI condition, the slope of the linear regression did not significantly differ from zero, suggesting no systematic change in accuracy across trials (see Table 2, Figure 8).

**Table 2:** Results of randomization test for the linear regression analysis of reaction times (RTs) and fraction of correct responses (FCRs) across trials. The table reports p-values derived from the permutation procedure, indicating whether the slope of the regression fitted to the original data differed significantly from slopes obtained from surrogate data. For each condition the original regression slope is provided. SLOW = slow breathing; FAST = fast breathing; NOI = no intervention; LISE = long inspiration/short expiration; SILE = short inspiration/long expiration.

|  | ***RT*** | ***FCR*** |
| --- | --- | --- |
| ***Slow*** |  |  |
| *Percentile* | 99.99 | 3.45 |
| *original slope* | 0.0064 | -0.007 |
| ***Fast*** |  |  |
| *Percentile* | 99.99 | 0.13 |
| *original slope* | 0.0064 | -0.011 |
| ***NOI*** |  |  |
| *Percentile* | 59.15 | 73.80 |
| *original slope* | 0.0012 | 0.0067 |
| ***SILE*** |  |  |
| *Percentile* | 100 | 6.35 |
| *original slope* | 0.018 | -0.01 |
| ***LISE*** |  |  |
| *Percentile* | 99.90 | 94.23 |
| *original slope* | 0.010 | 0.012 |

**Figure 7:**
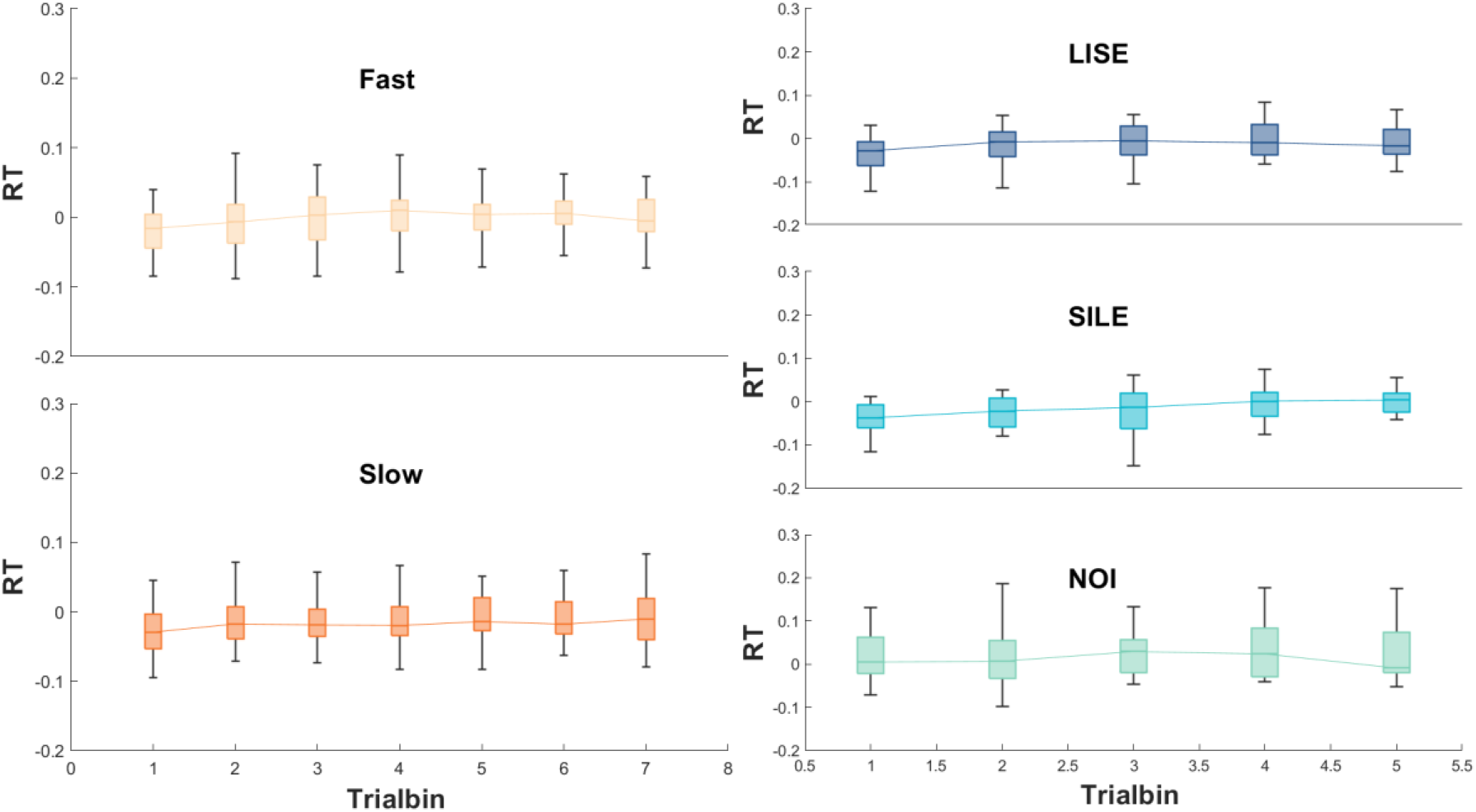
Temporal dynamics of RTs across trial bins for each breathing condition. The figure displays the binned reaction times (RTs) during the experimental blocks. Each panel shows the data for one breathing condition, with the two panels on the left corresponding to Experiment 1 (Slow and Fast breathing) and the three panels on the right to Experiment 2 (LISE, SILE, and NOI). For each participant, the RT per bin was computed as the mean RT of all trials within that bin after excluding trials based on criteria described in the Method section. Boxplots represent the distribution of normalized and squared RTs across participant. Results of the permutation tests assessing the significance of the linear slope across bins, as well as the original slope of the fitted regression line, are reported in Table 2.

**Figure 8:**
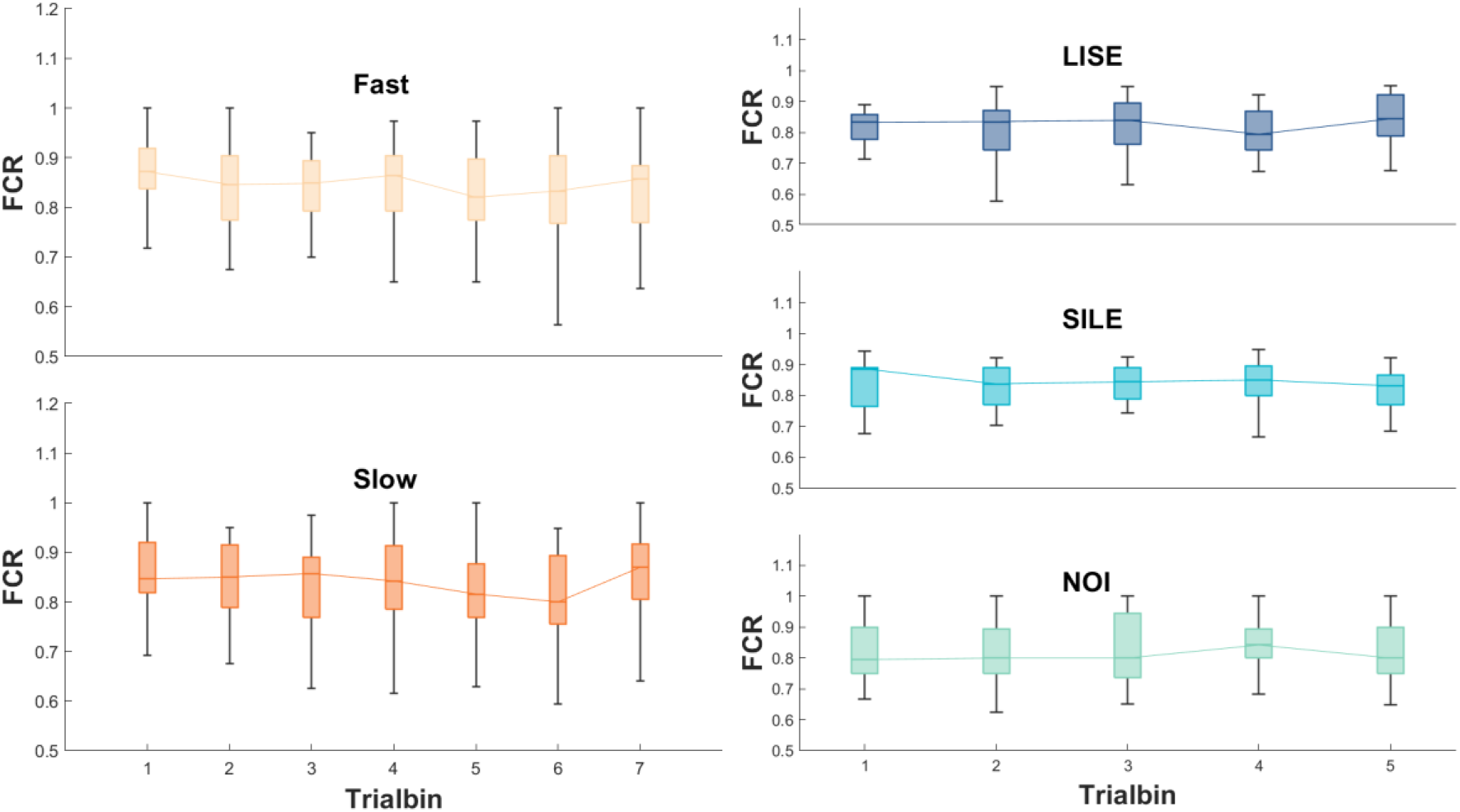
Temporal dynamics of task accuracy (FCR) across trial bins for each breathing condition. Figure illustrates changes in participants’ accuracy—indexed by the fraction of correct responses (FCR)—over the course of the post-practice task. Data are binned into segments of 10 consecutive trials, and for each participant. The FCR per bin was calculated as the proportion of correct responses among valid trials (see method section). The two panels on the left correspond to Experiment 1 (Slow and Fast breathing) and the three on the right representing Experiment 2 (LISE, SILE, and NOI). Boxplots show the distribution FCR values across participants. The outcome of the permutation-based slope analysis and the original slope values for each breathing condition are reported in Table 2.

## Discussion

### Adherence to guided breathing practice and post-practice respiratory dynamics

We studied the short-term effects of four guided breathing practices, probing how the application of a one-minute breathing practice affects participants respiratory dynamics following the practice and how this alters their performance in an emotion discrimination task.

Our results demonstrate that participants were generally able to adhere to instructed pattern of respiration. At the group-level these short-guided breathing protocols induced the expected changes in respiratory rate and inhalation to exhalation ratio. Particularly for the fast and slow breathing practice participants breathed significant faster (slower) than during spontaneous respiration, indicating the successful implementation of the externally paced respiration. For the patterns manipulating the inhalation-to-exhalation ratio we observed that the individual respiratory patterns during the breathing practice were more heterogeneous. Several participants exhibited I/E ratios opposing the intended pattern, highlighting potential difficulties in adhering to the instructed pattern. This highlights the importance of monitoring individual respiratory performance in studies investigating the impact of breathing practice.

When inspecting participants’ patterns of respiration in the minute immediately after the breathing practice, we observed an increased prevalence of atypical respiratory cycles compared to recordings of spontaneous respiration or conditions without a breathing practice. The emergence of such atypical cycles suggests that some participants’ required time to return to their typical respiratory pattern after the practice of conscious breathing and possibly followed the practice period with prolonged breath holds, sighs or other untypically structured respiratory cycles. We can only speculate about why this is the case. One possibility is that the application of respiration practice alters the physiological homeostatic balance (e.g. partial pressures of oxygen and carbon dioxide in the blood). The more this homeostatic balance is displaced from equilibrium, the stronger the compensatory response required to restore this to baseline. Alternatively, this reflects participants’ general flexibility in quickly adjusting their respiratory pattern. For sure, the individual variability in how accurately participants follow the instructed breathing practice, and how they breath following this requires more attention in future work.

The pattern of respiration following the breathing practice may also be critical for understanding what specific effects or how strong an effect of the specific practice actually is. Continuing to breathe similar to the instruction may amplify the effects, while a quick return to baseline may reduce them in comparison. The influence of atypical cycles, such as breath holds or sights, in turn remains unclear. This underscores the need to study participants breathing behavior more systematically during and following guided breathing interventions.

### Short-Lived Cognitive Effects of Breathing Practice and Potential Underlying Mechanisms

Despite inducing systematic changes in participants respiration, we did not find systematic differences in participants behavioral performance (FCR, or RTs) across the full post-practice blocks between the different breathing practices. Importantly, in experiment 2 we also did not see significant differences between the respiration conditions and a dedicated no-intervention manipulation. This speaks against longer-lived and systematic benefits following structured one-minute breathing practice, at least for the patterns and perceptual task studied here.

However, a time-resolved analysis revealed a transient improvement in reaction times immediately following the breathing practice. This suggests that even a brief period of structured breathing can enhance task responsiveness, although the benefit appears short-lived and dissipates over the time scale of tens of seconds. One interpretation of the transient enhancement in RTs is a short-term shift in autonomic state. Although an influence via the hypothalamic–pituitary–adrenal (HPA) axis appears unlikely on the short time scale considered here a potential involvement of this system cannot be entirely ruled out, given that studies on long term practice of deep breathing have demonstrated systematic changes in cortisol levels (Ma et al. 2017; Martarelli et al. 2011; Perciavalle et al. 2017). Measuring cortisol concentrations in saliva immediately following different breathing practice could clarify whether distinct patterns of HPA axis activation underlie observed cognitive effects. An alternative account is the involvement of changes in the balance of the autonomic nervous system. Previous work has shown that slow-paced breathing (De Couck et al. 2019), hyperventilation (Neginhal et al. 2017), and modified I/E ratio patterns (van Diest et al. 2014) lead to changes in the heart rate variability (HRV). Given the well-established link between vagal tone and cognitive control as proposed by the neurovisceral integration model (Smith et al. 2017; Thayer und Lane 2000; Thayer und Lane 2009) a change in HRV caused by respiration practice may reflect changes in the autonomic nervous system through which breathing practice could indirectly modulates cognitive performance. Furthermore, the gradual increase in RTs over tens of trials may reflect a return to physiological baseline, consistent with a homeostatic reset in cardiovascular or neural parameters. If true, one would expect more pronounced and lasting improvements in cognition following long-term breathing practice that leads to a sustained enhancements in HRV (Balban et al. 2023).

Alternatively, or possibly complementary, respiration may directly modulate large-scale brain activity. Recent studies have demonstrated that respiration and neural oscillations are tightly coupled during both rest and task engagement (Kluger und Gross 2020; Kluger et al. 2021; Zelano et al. 2016). These respiration-locked modulations in oscillatory power have been shown to account for variations in task performance, suggesting that rhythmic changes in neural excitability may underlie respiration-related fluctuations in perception and cognitive processing ((Kluger et al. 2021). Such modulations of neural oscitations have also been observed during high-frequency yoga breathing (Budhi et al. 2024), voluntary deep breathing (Karjalainen et al. 2025) and slow controlled breathing (Hsu et al. 2020). Transient modulations of cortical dynamics may provide a mechanistic explanation for the short-lived behavioral improvements in reaction times observed in the present study. Once the breathing practice ends and spontaneous respiration resumes, the induced neural state likely returns to baseline, which could account for the temporal limitation of the observed effects. However, there is currently little evidence regarding how different breathing practices, especially those manipulating the inhalation-to-exhalation ratio, affect oscillatory activity, representing an important open question for future research

### Limitations and outlook

The present study has clear limitations. First, the duration of the breathing practice was short (one minute), which may not be sufficient to induce lasting and sustained changes in bodily physiology or the regulation of the autonomous nervous system. This short duration may explain the limited differentiation in effects in the cognitive task across techniques. Second, although we analyzed individual respiratory traces, we did not systematically monitor the physiological parameters such as heart rate, heart rate variability or respiratory gas composition (e.g., CO₂ or O₂ concentration). This makes it difficult to link the individual variability in respiratory patterns to the outcome. Third, the no intervention condition, in which participants sat quietly and breathed naturally, served as a reference for spontaneous respiration and physiological resting state rather than controlling for attentional engagement or task anticipation. It allowed comparison against structured breathing practice to ensure that observed effects were not driven by natural variability or temporal factors. Future studies should implement control conditions such as breath awareness or heartbeat counting tasks to isolate the effects of attentional focus on interoceptive signals from those attributable to the alteration of breathing patterns per se. Finally, we did not measure brain activity (e.g. by using EEG) during or after the breathing practice. This limits our ability to link respiration-induced behavioral changes to underlying neurophysiological mechanisms. Addressing these points in future research will be essential for more precisely disentangling the pathways through which respiratory practice influences cognitive function.

## Conclusion

The present study demonstrates that even brief applications of guided breathing practices can transiently modulate cognitive performance, as reflected in faster reaction times immediately following the practice. While the effects were short-lived and may be independent of the specific breathing practice, they suggest a general facilitative impact of structured respiration on attentional readiness or sensorimotor responsiveness. These findings underscore the feasibility of implementing guided breathing in experimental paradigms, highlight the relevance of temporal dynamics when assessing their cognitive effects and raise important questions about the underlying mechanisms.

